# CREB-pCREB-PER2 feedback loop mediates transition between mania and depression-like behaviors

**DOI:** 10.1101/2022.09.28.509998

**Authors:** Xin-Ling Wang

## Abstract

Little is known about the mechanisms underlying the transition between mania and depression. We found here that ouabain decreased pCREB and PER2 levels in CA1 and induced mania-like behavior, which was attenuated by lithium and overexpression of *Per2* in this region. Furthermore, knockdown of *Per2* in CA1 induced mania-like behavior, in contrast, overexpression of *Per2* resulted in depression-like behavior. Similar results were found in manipulations of *Creb1* in CA1. Western blot analyses revealed that upregulations of CREB or PER2 can increase each other’s levels, besides pCREB, and vice versa. Therefore, the CREB– pCREB–PER2 pathway forms a positive feedback loop that mediates the transition between manic and depressive phenotypes.

**One-Sentence Summary:** A novel molecular loop underlies phase inversion of bipolar disorder

## Main Text

Bipolar disorder (BD) is a severe psychiatric disorder characterized by alternating manic/hypomanic episodes with depressive episodes. BD is the 17th most common disability globally (*1*). The World Mental Health Survey reported that the lifetime prevalence and 12-month prevalence of BD were 2.4% and 1.5%, respectively (*2*). Throughout the course of BD, the manic/hypomanic, depressive episodes and symptom-free periods can transit mutually. Notably, some antidepressants have the risk of inducing manic episodes in BD patients (*3, 4*). Therefore, preventing or modulating the switch between manic and depressive episodes may have great therapeutic potential for BD. A large number of studies have reported that circadian rhythm and clock genes play critical roles in mood regulation in BD (*5-13*). However, because of the limitation of animal models (*14, 15*), the mechanism underlying phase transitions in BD remains unclear. Young et al. reported that mice with reduced dopamine transporter levels exhibited seasonal change-induced switching between states similar to BD (*16*). However, this model does not adequately mimic the pathophysiology of BD (*17*).

The *Period* family of clock genes include three homologous genes—*Per1, Per2* and *Per3*. There have been numerous reports on the involvement of *Per1* and *Per2* in affective disorders (*18*). A recent study reported that *Per1* knockout mice exhibited depression-like behaviors, which could be ameliorated by upregulation of *Per1* in the lateral stiff nucleus (*19*). Knockout of *Per2* in glial cells also attenuated depression-like behavior in mice (*20*). There have been also many reports on the involvement of *Per2* in the pathogenesis of BD (*21-24*). Through pharmacogenetics, Moreira *et al*. identified several clock genes associated with the response to lithium, including *NR1D1, GSK3beta, CRY1, ARNTL, TIM* and *PER2* (*25*). Li *et al*. demonstrated that lithium increased *Per2* expression at both transcriptional and post-transcriptional levels (*26*). Furthermore, Kim and his colleagues found that lithium upregulated *Per2* and *ERG1* (*22*). A recent study by Zhou *et al*. showed that lithium increased *Per2* expression by decreasing the expression of the transcription factor E4BP4 (*23*). Taken together, these previous studies suggest a key role of *Per2* in the pathogenesis of BD. However, the molecular mechanisms underlying BD and phase transition are still unknown.

cAMP response element-binding protein (CREB) 1 is a transcription factor that regulates hundreds of genes containing CREB binding sites (*27, 28*), and can be phosphorylated at Ser133 by a variety of signals, including cyclic adenosine monophosphate (cAMP), calcium, MAPK, PKA, PKC and stress signals (*27-30*). Phosphorylated CREB (pCREB), a known transcriptional regulator of *Per2*, has also been reported to be involved in the pathogenesis of BD (*31-33*). For example, Gaspar *et al*. showed that elevated levels of CREB/pCREB are found in cells from patients with BD (*34*). Furthermore, clinical studies showed that pCREB levels are significantly increased in the lymphoblasts from peripheral blood of patients with BD (*35*). However, most of these were invitro studies, and did not reveal how pCREB-*Per2* pathway alters upon phase transition in invivo studies.

In this study, we aim to investigate how CREB/pCREB-*Per2* pathway acts on mood regulation. We found that CREB/pCREB positively regulates *Per2* expression, which also positively regulates CREB/pCREB in a feedback way, forming a loop in CA1. Besides, downregulation of this loop in CA1 produces mania-like behavior, while upregulation of it in this region leads to depression-like behavior.

### pCREB and PER2 in the CA1 are involved in ouabain-induced mania-like behavior

To investigate whether pCREB and PER2 play roles in the pathogenesis of BD, we produced the ouabain-induced model of BD by infusing ouabain dissolved in artificial cerebrospinal fluid (aCSF) (1 mmol/L, 5 μL) intracerebroventricularly (ICV) in rats (Fig. 1, A and B and fig. S1A), as described in previous studies (*31*). Rats in the control group were infused with aCSF in the same region (Fig. 1B). Then, we administered vehicle (saline) or lithium carbonate (20 mg/kg), the treatment for BD, intraperitoneally into rats once per day for 7 consecutive days (Fig. 1A and fig. S1A) (*31*). We used the sucrose preference test (SPT), forced swim test (FST), open field test (OPT) and elevated plus maze (EPM) test to assess behaviors (*36*). On day 5 of administration, sucrose preference values were significantly increased in the ouabain group compared with the control group (fig. S1B), indicating a mania-like behavior, which was significantly ameliorated by lithium carbonate (fig. S1B). On day 7, in the FST, the immobility time was significantly reduced in the ouabain group compared with the control group (fig. S1C), and this was significantly ameliorated by lithium (fig. S1C), demonstrating an anti-mania-like effect of lithium. In the OPT, total distance in the ouabain group was significantly increased compared with the control group (fig. S1, D and E), indicating a hyperactivity behavior, which was significantly mitigated in the ouabain + lithium group (fig. S1, D and E). In parallel, time in central zones was significantly increased in the ouabain group compared with the control group (fig. S1, D and F), indicating an increased exploratory behavior. Together, these results show that ouabain successfully models mania, which can be attenuated by lithium, in line with previous studies (*36*).

To explore the molecular changes underlying these behaviors, we conducted western blot analysis in proteins extracted from CA1 region, which was shown to play great roles in mood regulation (*37*). Results showed that PER2 and pCREB levels in the CA1 region were significantly decreased in the ouabain group compared with the control group (Fig. 1, C-E), and these reductions were significantly ameliorated by lithium (Fig. 1, D and E), in line with previous studies (*22, 23, 29, 31, 35, 38, 39*). There were no significant differences in CREB between groups (Fig. 1F). Thus, pCREB and PER2 in the hippocampal CA1 may play critical roles in mania-like behavior.

**Fig. 1.**
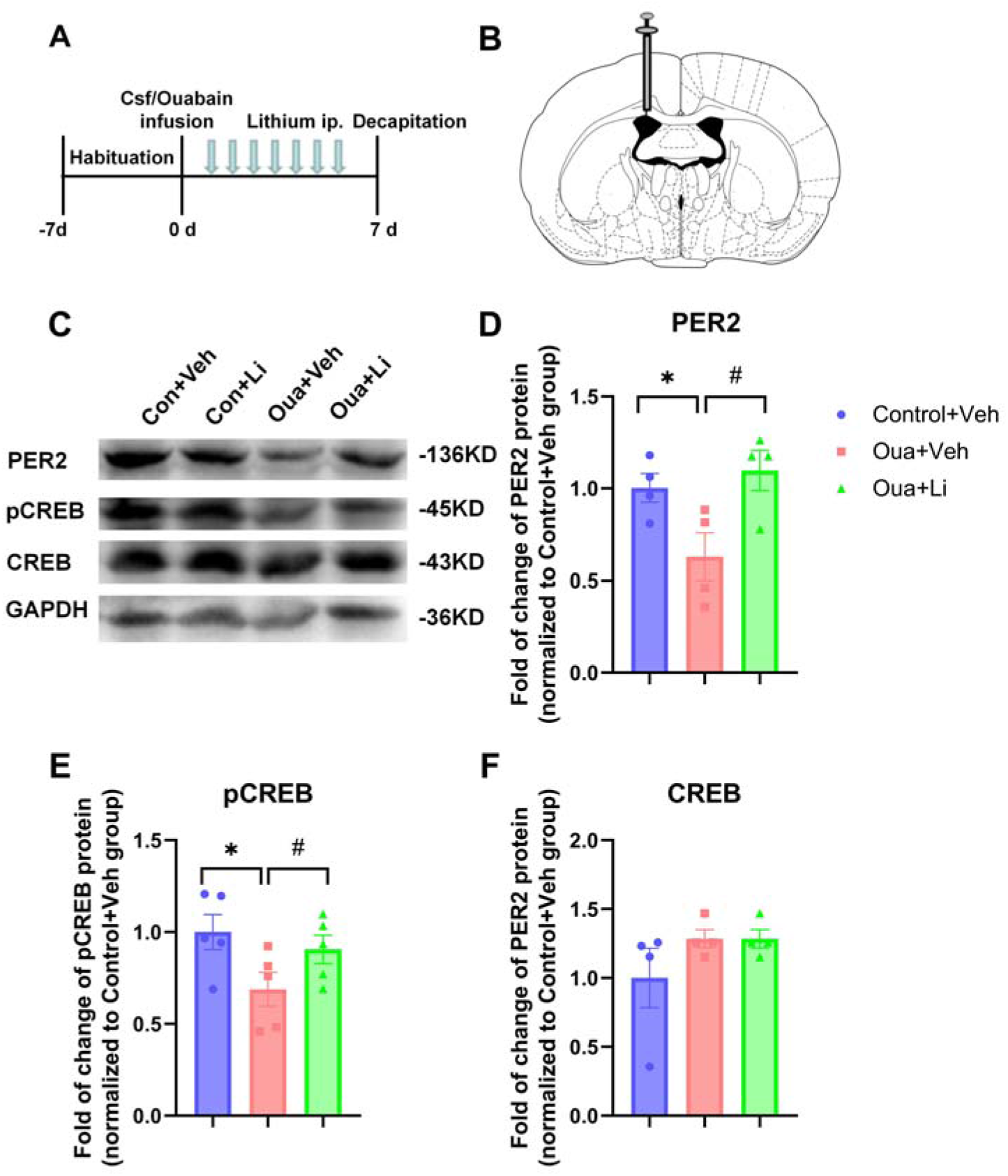
pCREB and PER2 in CA1 were potentially involved in mania-like behavior. (**A**)Timeline of the experiment. (**B**) Schematic of the infusion site of aCSF or ouabain in the rat brain: anterior/posterior, -0.9 mm; medial/lateral, + 1.5 mm; and dorsal/ventral, -3.3 mm. Modified from the rat brain atlas. (**C**) Representative western blots reflecting changes in expression of PER2, pCREB and CREB protein levels in the control and ouabain administered vehicle or lithium groups. Molecular weights are indicated on the right. (**D** to **F**) Quantification of the relative protein levels of PER2 (D), pCREB (E) and CREB (F). All data are presented as mean ± SEM. *n* = 4-5, values exceeded mean ± 2 * STD were excluded. \**P* < 0.05, vs. control + vehicle group; #*P* < 0.05, vs. ouabain + vehicle group; two-tailed unpaired *t* test.

### Knockdown of *Per2* in CA1 produces mania-like behavior and downregulates CREB levels

To investigate the role of *Per2* in CA1 region in mood regulation, we performed knockdown of *Per2* in this region and conducted behavioral tests (Fig. 2A). After rats were habituated for 7 days, we microinjected adeno-associated virus (AAV)-sh*Per2* into CA1 region to downregulate expression of *Per2*. Then, 21 days later, we examined protein levels in CA1 (fig. S2A) and performed behavioral tests, including the SPT, FST, OPT and EPM test (Fig. 2A). In SPT, sucrose preference values were significantly increased in the AAV-sh*Per2* group, compared with the control group (Fig. 2C). Furthermore, immobility time in the *Per2*-knockdown group was significantly shortened vs. the control group (Fig. 2D), indicating a mania-like behavior. In EPM test, time spent in open arms was significantly increased in the AAV-sh*Per2* group, compared with the control group (Fig. 2, E and G). There was no significant difference between the two groups in number of entries into open arms (Fig. 2, E and F). In OPT, total distance traveled was significantly increased in the AAV-sh*Per2* group, compared with the control group (Fig. 2, H and I), indicating a hyperactivity behavior. Moreover, time in central zones was also significantly increased in this group (Fig. 2, H and J), demonstrating an increased exploratory behavior, a mania-like behavior. Collectively, these behavioral results demonstrate that knockdown of *Per2* in CA1 causes mania-like behaviors in rats.

**Fig. 2.**
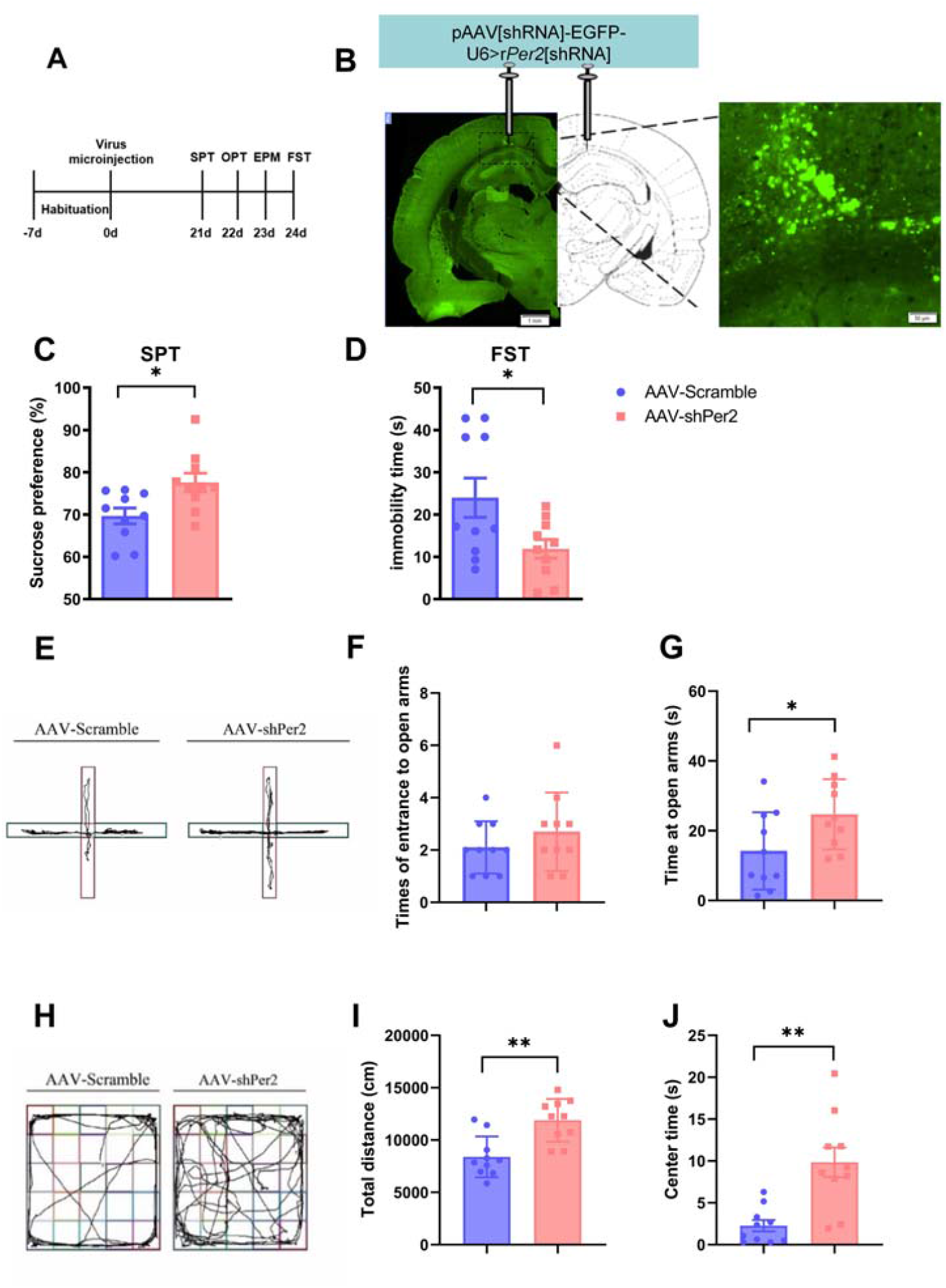
Knockdown of *Per2* in CA1 region produced mania-like behavior. (**A**) Timeline of the experiment. (**B**) Representative immunofluorescence images showing injection sites of AAV (left panel, under 10 * object lens) and transfection cells (in fluorescence green, right panel, under 40 * object lens) in CA1 of the rat brain. Scale bars, 1 mm (left), 50 μm (right). (**C** and **D**) Data show the effects of AAV-shPer2 infused into CA1 on sucrose preference values in SPT (C) and immobility time in FST (D). Graphs are shown as means ± SEMs. *n* = 10. \**P* < 0.05; unpaired *t* test. (**E** to **G**) Results of EPM test in AAV-scramble and AAV-shPer2 groups. (E) shows representative images of rat traces of both groups on the elevated plus maze. Graphs are times of entries to open arms (F) and time spent at open arms (G) in both groups. Data are shown as means ± SEMs. *n* = 10. \**P* < 0.05; unpaired *t* test. (**H** to **J**) Open field test reults: (H) Representative images of rat traces of the two groups in open field test. Graphs are total distance (I) and time spent in central zones (J) in both groups. Data are shown as means ± SEMs. *n* = 10. \*\**P* < 0.01; unpaired *t* test (I), Mann Whitney test (J).

Western blot analysis validated that PER2 levels were significantly downregulated by AAV-sh*Per2* (fig. S2C). Besides, CREB levels in CA1 were also significantly reduced in the AAV-sh*Per2* group, compared with the AAV-scramble group (fig. S2E). Therefore, knockdown of *Per2* in CA1 also downregulates CREB in this brain region.

### Overexpression of *Per2* in CA1 induces depression-like behavior, ameliorates ouabain-induced mania-like behavior and upregulates CREB/pCREB levels in this region

To further examine the function of PER2 in CA1 in mood modulation, we overexpressed *Per2* in the CA1 by microinjection of lentivirus (LV)-*Per2* into this region, and performed behavioral tests (Fig. 3A). Sucrose preference values were significantly reduced in the LV-*Per2* group, compared with those in the control group (Fig. 3C), an indicative of depression-like behavior. In FST, immobility time was significantly prolonged in the LV-*Per2* group, also indicating a depression-like behavior (Fig. 3D). Moreover, in the EPM and OPT, time in open arms as well as time in central zones were significantly reduced in the LV-*Per2* group vs. the control group (Fig. 3, G and J), indicative of anxiety-like behaviors. Therefore, overexpression of *Per2* in CA1 induces depression-like behaviors.

**Fig. 3.**
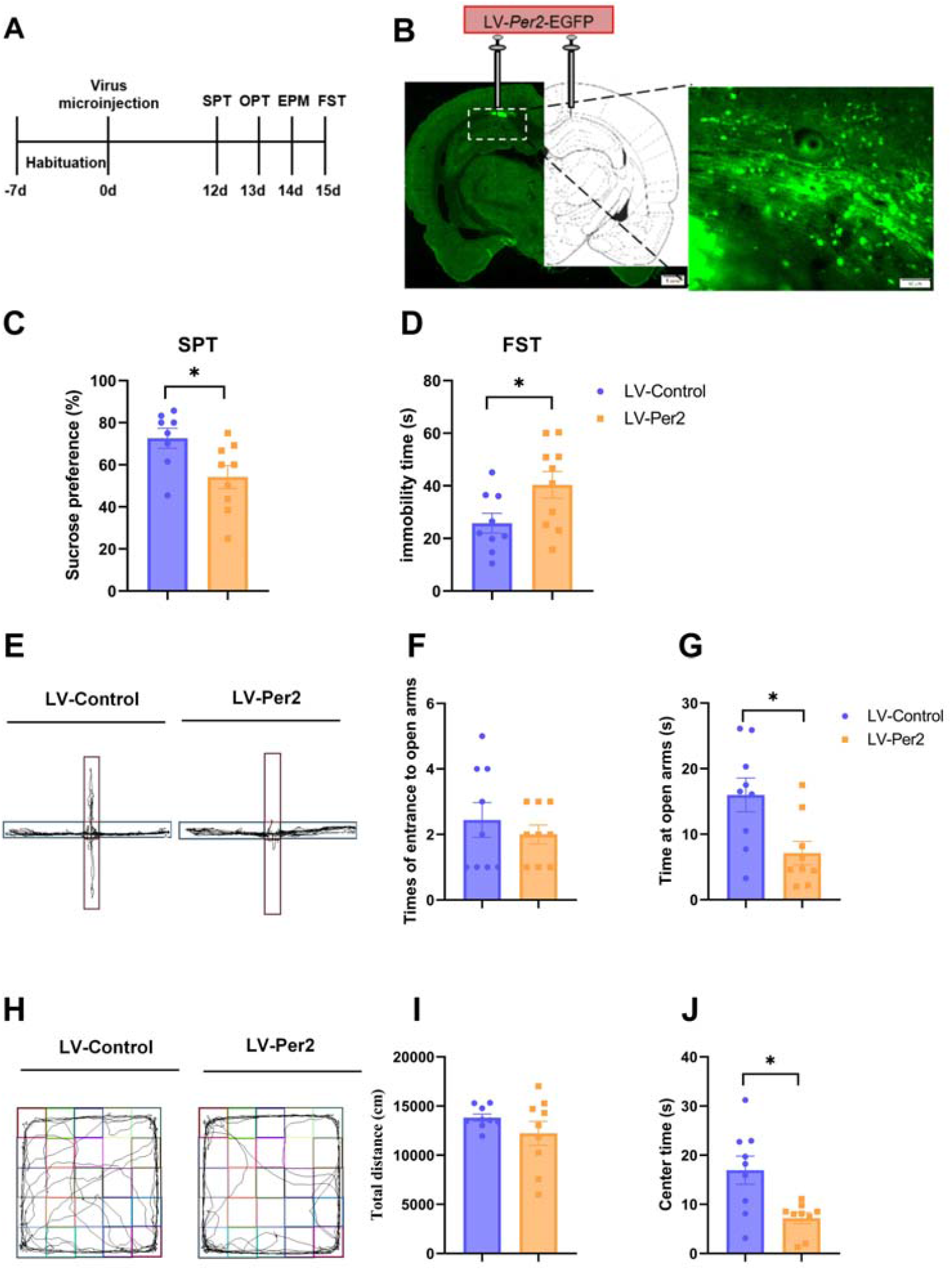
Overexpression of *Per2* in CA1 region induced depression-like behavior. (**A**) Timeline of the experiment. (**B**) Representative immunofluorescence images showing injection sites of LV (left panel, under 10 * object lens) and transfection cells (in fluorescence green, right panel, under 40 * object lens) in CA1 of the rat brain. Scale bars, 1 mm (left), 50 μm (right). (**C** and **D**) Data showing the effects of LV-Per2 infused into CA1 on sucrose preference values in SPT (C) and immobility time in FST (D). Graphs are shown as means ± SEMs. *n* = 8-10, values exceeded mean ± 2 * STD were excluded. \**P* < 0.05; unpaired *t* test. (**E** to **G**) Results of EPM in LV-control and LV-Per2 group: representative images of rat traces of both groups on the elevated plus maze (E), times of entrance to open arms (F) and time spent at open arms (G). Bar charts are shown as means ± SEMs. *n* = 9. \**P* < 0.05; unpaired *t* test. (**H** to **J**) Results of OPT in LV-control and LV-Per2 group: representative images of rat traces of the two groups in the open field (H), total distance (I) and time spent in central zones (J). Bar charts are shown as means ± SEMs. *n* = 9. \**P* < 0.05; Mann Whitney test.

To confirm the role of *Per2* in the pathogenesis of BD, we also infused ouabain into ICV to model BD and microinjected LV-*Per2* into the CA1 simultaneously to examine whether it can improve ouabain-induced manic behavior. Behavioral tests were performed 5–7 days after injection of CSF/ouabain and lentivirus, including SPT, OPT, EPM and FST (fig. S4A). In SPT, there were no significant differences in sucrose preference between groups (fig. S4B). In FST, immobility time was significantly higher in the ouabain + LV-control group, compared with the control group (fig. S4C), indicating a mania-like behavior, which was significantly attenuated by LV-*Per2* (fig. S4C), indicating an anti-manic effect of LV-*Per2* in CA1. In EPM, time in open arms was significantly prolonged by ouabain, which was significantly shortened by LV-*Per2* (fig. S4, D and F), also demonstrating an anti-mania like action. In OPT, time spent in central zones was significantly increased in the ouabain + LV-control group, which was significantly reduced by LV-*Per2* (fig. S4, G and I), also indicating an anti-manic like effect of LV-*Per2* in CA1.

Western blot analysis of proteins in CA1 was performed 12 days after lentivirus injection (fig. S3A). PER2 levels were verified to be significantly upregulated in CA1 (fig. S3C), furthermore, pCREB and CREB levels were also significantly elevated in the LV-*Per2* group (fig. S3, D and E), Thus, overexpression of *Per2* in CA1 upregulates CREB and pCREB in this region.

### Knockdown of CREB in CA1 induces mania-like behavior, which can be relieved by overexpression of *Per2* in this region

To explore the role of CREB, an upstream molecule of *Per2*, in mood-related behaviors and its interaction with *Per2*, and performed knockdown of *CREB1* by infusing AAV-sh*CREB1* bilaterally into the CA1 (Fig. 4B), followed by behavioral tests 21 days later (Fig. 4A). Notably, sucrose preference values were significantly increased and immobility time was significantly reduced in the AAV-sh*CREB* group, compared with the control group (Fig. 4, C and D), suggestive of mania-like behaviors. In the EPM test, number of entries into open arms and time spent in the open arms were both significantly increased in the *CREB* knockdown group vs. the control group (Fig. 4, E–G). Furthermore, total distance and time spent in central zones were both significantly increased in this group vs. the control group (Fig. 4, H–J), also exhibiting mania-like behaviors. Next, we infused LV-*Per2* into CA1 bilaterally to examine whether these behaviors can be ameliorated by overexpressed PER2. Behavioral tests were performed 12 days after injection (Fig. 4A). Sucrose preference values and immobility time were significantly improved in the AAV-sh*CREB* group post injection compared with those before injection (Fig. 4, C and D). In the EPM test, entries into open arms and time at open arms were both decreased significantly by LV-*Per2* in this group vs. those before virus injection (Fig. 4, E–G), demonstrating an anti-manic like effect of overexpressed PER2 in CA1. In OPT, time in central zones was also moderated by injection of LV-*Per2* (Fig. 4, H and J).

**Fig. 4.**
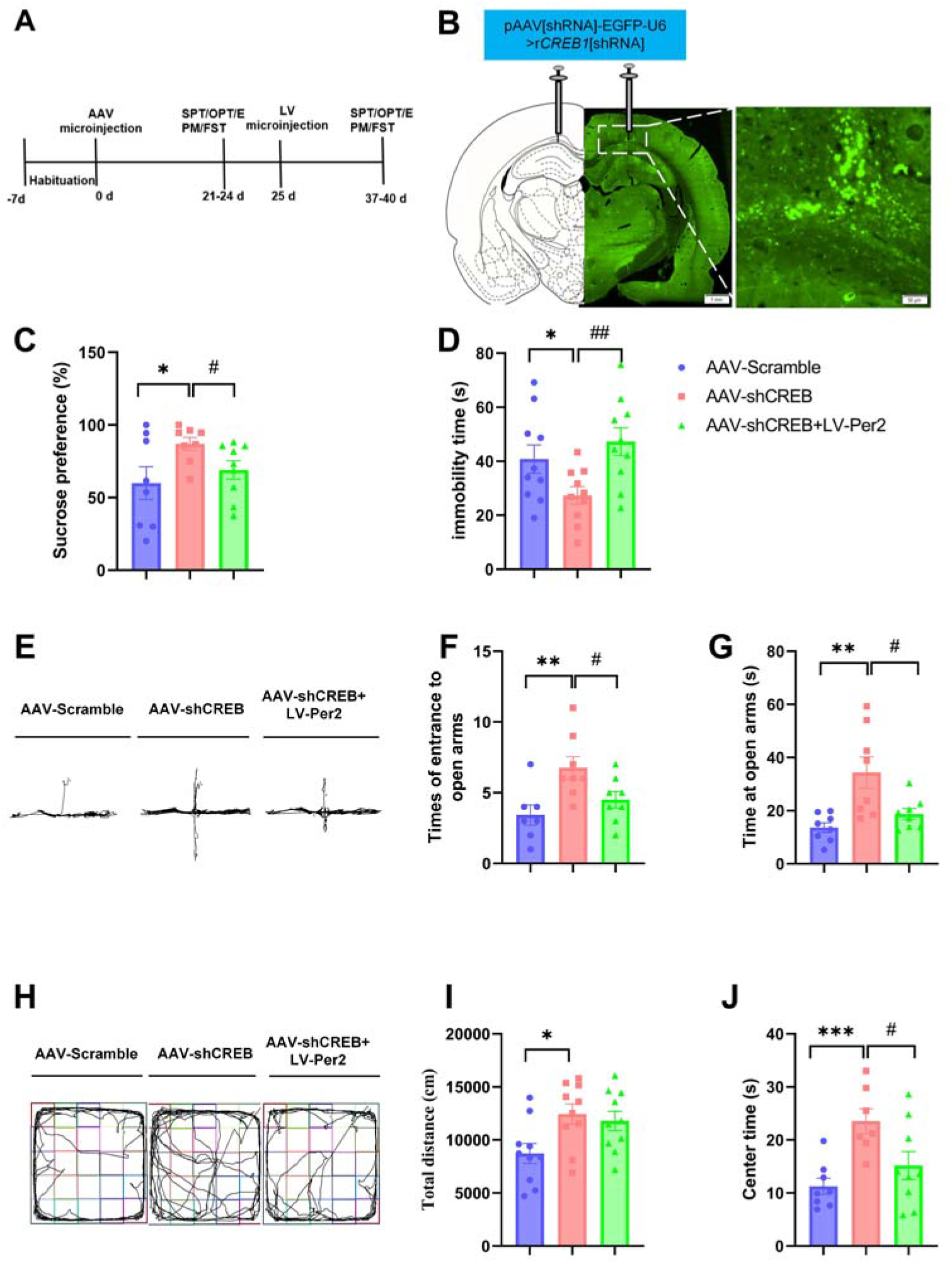
Knockdown of *CREB* in CA1 produced mania-like behaviors, which can be rescued by overexpression of *Per2* in this region. (**A**) Timeline of this experiment. (**B**) Representative immunofluorescence images showing injection sites of AAV (left panel, under 10 * object lens) and transfection cells (in fluorescence green, right panel, under 40 * object lens) in CA1 of the rat brain. Scale bars, 1 mm (left), 50 μm (right). (**C** and **D**) Data showing the effects of AAV-shCREB and LV-Per2 infused into CA1 on sucrose preference values in SPT (C) and immobility time in FST (D). Graphs are shown as means ± SEMs. *n* = 8-10, values exceeded mean ± 2 * STD were excluded. \**P* < 0.05, vs. AAV-scramble group; #*P* < 0.05, ##*P* < 0.01, vs. AAV-shCREB group; unpaired *t* test. (**E** to **G**) Results of EPM in AAV-scramble, AAV-shCREB and AAV-shCREB + LV-Per2 groups: representative images of rat traces of the three groups on the elevated plus maze (E), times of entrance to open arms (F) and time spent at open arms (G). Bar Graphs are shown as means ± SEMs. *n* = 7-8. \*\**P* < 0.01, vs. AAV-scramble group; #*P* < 0.05, vs. AAV-shCREB group; unpaired *t* test. #*P* < 0.05, Mann Whitney test (G). (**H** to **J**) Results of OPT: representative images of rat traces in the open field (H), total distance (I) and time spent in central zones (J). Bar Graphs are shown as means ± SEMs. *n* = 7-10. \**P* < 0.05, \*\*\**P* < 0.001, vs. AAV-scramble group; #*P* < 0.05, vs. the AAV-shCREB group; unpaired *t* test.

Western blot analysis validated that CREB were downregulated significantly by AAV-shCREB, and can be relieved by LV-*Per2* (fig. S5E). Besides, PER2 and pCREB levels were also decreased significantly by AAV-sh*CREB* (fig. S5, B-D). Therefore, knockdown of CREB can also downregulates pCREB and PER2, and changes in CREB can be rescued by overexpression of *Per2*. In line with our results, McCarthy et al. also demonstrated that knockdown of CREB in fibroblasts from BD patients leads to a reduction in the baseline amplitude of PER2 rhythmicity (*38*).

### Overexpression of CREB in CA1 causes depression-like behavior, which can be relieved or converted to mania-like behavior by knockdown of *CREB1* or *Per2* in this region

To further investigate the role of CREB and its crosstalk with PER2 in mood-related behaviors, we overexpressed *CREB1* by infusing LV-*CREB*, and at the same time, we injected AAV-scramble, AAV-sh*CREB* or AAV-sh*Per2* into CA1 bilaterally (Fig. 5, A and B), and conducted behavioral tests 21 days later (Fig. 5A). The sucrose preference values were significantly decreased and immobility time was significantly increased in the LV-*CREB* + AAV-scramble group, demonstrating depression-like behaviors (Fig. 5, C and D), in line with previous reports (*27*). Moreover, sucrose preference values that were reduced by LV-*CREB* can be significantly countered by AAV-sh*CREB* or AAV-sh*Per2* (Fig. 5C), besides, the immobility time was also improved significantly by AAV-sh*Per2* (Fig. 5D). Surprisingly, sucrose preference values were even higher in the LV-*CREB* + AAV-sh*Per2* group vs. the control group (Fig. 5C), indicating a probable transitioning to mania-like behavior by AAV-sh*Per2*. In OPT, total distance and time in central zones were both reduced significantly by LV-*CREB*, indicating anxiety-like behaviors (Fig. 5, E and F), which could be diminished by AAV-sh*CREB* or AAV-sh*Per2* (Fig. 5, E and F). In the EPM test, there was no significant difference in number of entries into open arms in either LV-*CREB* + AAV-sh*CREB* or LV-*CREB* + AAV-sh*Per2* group vs. the control group (Fig. 5G), indicating the anxiolytic-like effects of AAV-sh*CREB* and AAV-sh*Per2*. However, time at open arms was still significantly lower in the LV-*CREB* + AAV-sh*Per2* group, compared vs. the control group, showing a limited anxiolytic-like effect of AAV-sh*Per2* (Fig. 5H).

**Fig. 5.**
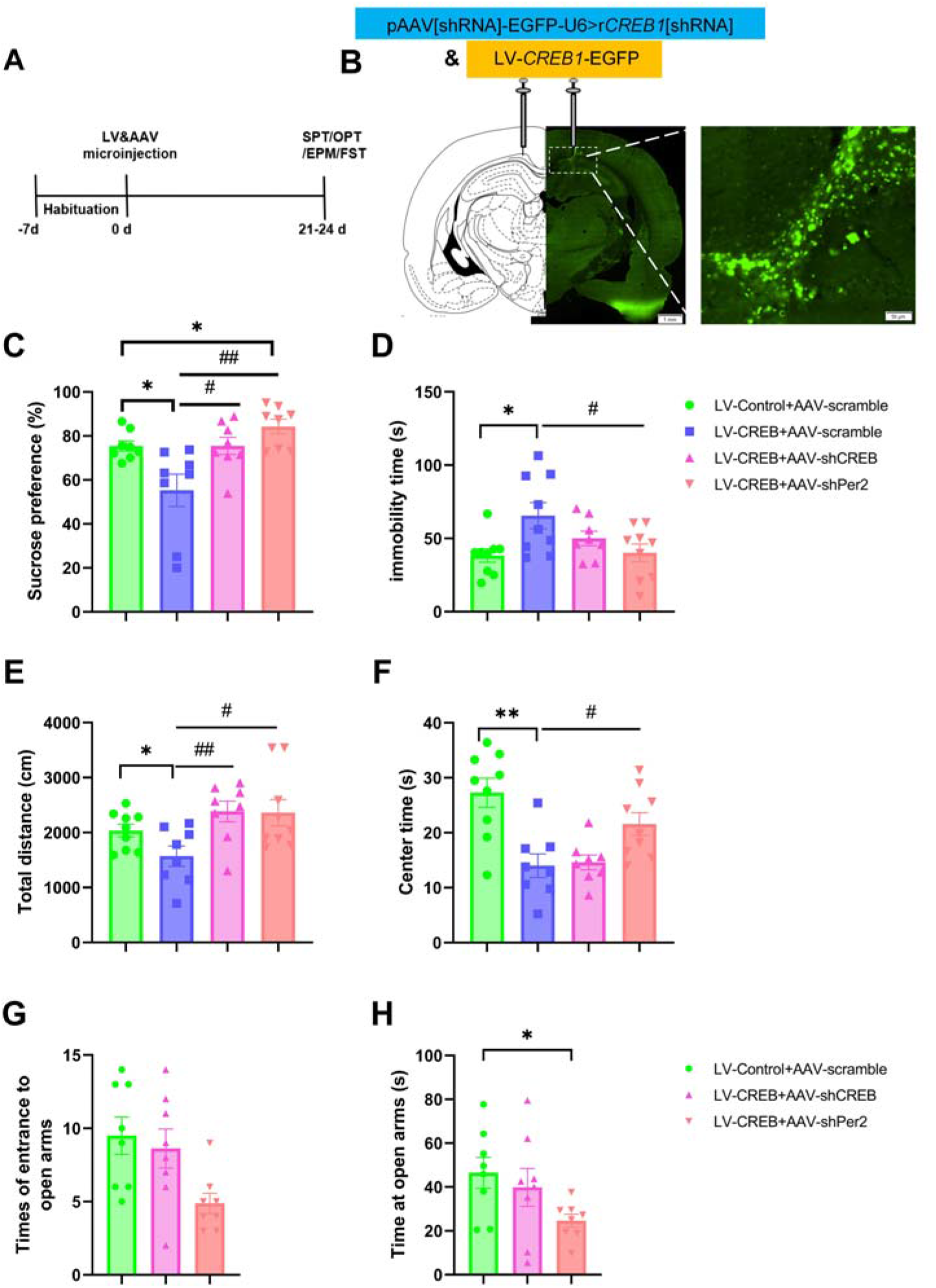
Overexpression of *CREB1* in CA1 induced depression-like behaviors, which can be mitigated or converted by knockdown of either *CREB1* or *Per2* in this region. (**A**) Timeline of this experiment. (**B**) Representative immunofluorescence images showing coinjection sites of LV and AAV (left panel, under 10 * object lens) and transfection cells (in fluorescence green, right panel, under 40 * object lens) in CA1. Scale bars, 1 mm (left), 50 μm (right). (**C** and **D**) Data showing the effects of LV-CREB and AAV-shCREB/shPer2 infused into CA1 on sucrose preference values in SPT (C) and immobility time in FST (D). Graphs are shown as means ± SEMs. *n* = 8-9, values exceeded mean ± 2 * STD were excluded. \**P* < 0.05, vs. LV-control + AAV-scramble group; #*P* < 0.05, ##*P* < 0.01, vs. LV-shCREB + AAV-scramble group; unpaired *t* test. \**P* < 0.05, Mann Whitney test (C). (**E** and **F**) Results of OPT: total distance (E) and time spent in central zones (F) in the four groups. Graphs are shown as means ± SEMs. *n* = 8-9. \**P* < 0.05, \*\**P* < 0.01, vs. LV-control + AAV-scramble group; #*P* < 0.05, ##*P* < 0.01, vs. LV-CREB + AAV-scramble group; unpaired *t* test. (**G** and **H**) Results of EPM: times of entries to open arms (G) and time spent at open arms (H). Graphs are shown as means ± SEMs. *n* = 8. \**P* < 0.05, vs. LV-control + AAV-scramble group; Mann Whitney test (H).

Western blotting of protein extracted from rat CA1 regions in LV-control and LV-*CREB* groups validated that CREB was significantly upregulated by LV-*CREB* (fig. S6E). PER2 and pCREB levels were also significantly upregulated by LV-*CREB* (fig. S6, C and D), indicating a positive regulatory effect of CREB on pCREB and PER2.

### Mania-like behaviors induced by knockdown of *Per2* in CA1 can be ameliorated or converted by overexpression of *Per2* or *CREB1* in this region

As shown in Fig. 2, knockdown of *Per2* in CA1 produced mania-like behaviors. To examine whether these behaviors can be ameliorated or converted by overexpression of *Per2* or *CREB1* in CA1, we microinjected AAV-sh*Per2* bilaterally into this region on day 1, followed by infusing LV-*Per2* or LV-*CREB* 9 days later, and performing behavioral tests another 12 days later (Fig. 6A). Representative infusion sites of AAV and LV in CA1 are shown in Fig. 6B. Sucrose preference values were significantly increased in the AAV-sh*Per2* group vs. the control group, nevertheless, they were not significantly altered in the AAV-sh*Per2* + LV-*Per2* or AAV-sh*Per2* + LV-*CREB* group vs. the AAV-sh*Per2* group (Fig. 6C). While, in FST, inconsistent with former results in Fig. 2, immobility time was significantly reduced in the AAV-sh*Per2* group vs. the control group, and this was significantly altered by LV-*Per2* or LV-*CREB* (Fig. 6D), revealing anti-manic like effects of overexpressed PER2 and CREB in CA1. Furthermore, in OPT, both total distance and time in central zones were significantly increased by AAV-sh*Per2*, in line with results above (Fig. 2, I and J), and these were suppressed by LV-*Per2* or LV-*CREB* (Fig. 6, E and F), further supporting an anti-manic like action of overexpressed PER2 and CREB. Furthermore, in the EPM test, times of entries into open arms showed no significant difference among different groups (Fig. 6G), while, time at open arms was significantly decreased in the AAV-sh*Per2* + LV-*CREB* group vs. the control group (Fig. 6H), suggesting a potential conversion to anxiety-like behavior.

**Fig. 6.**
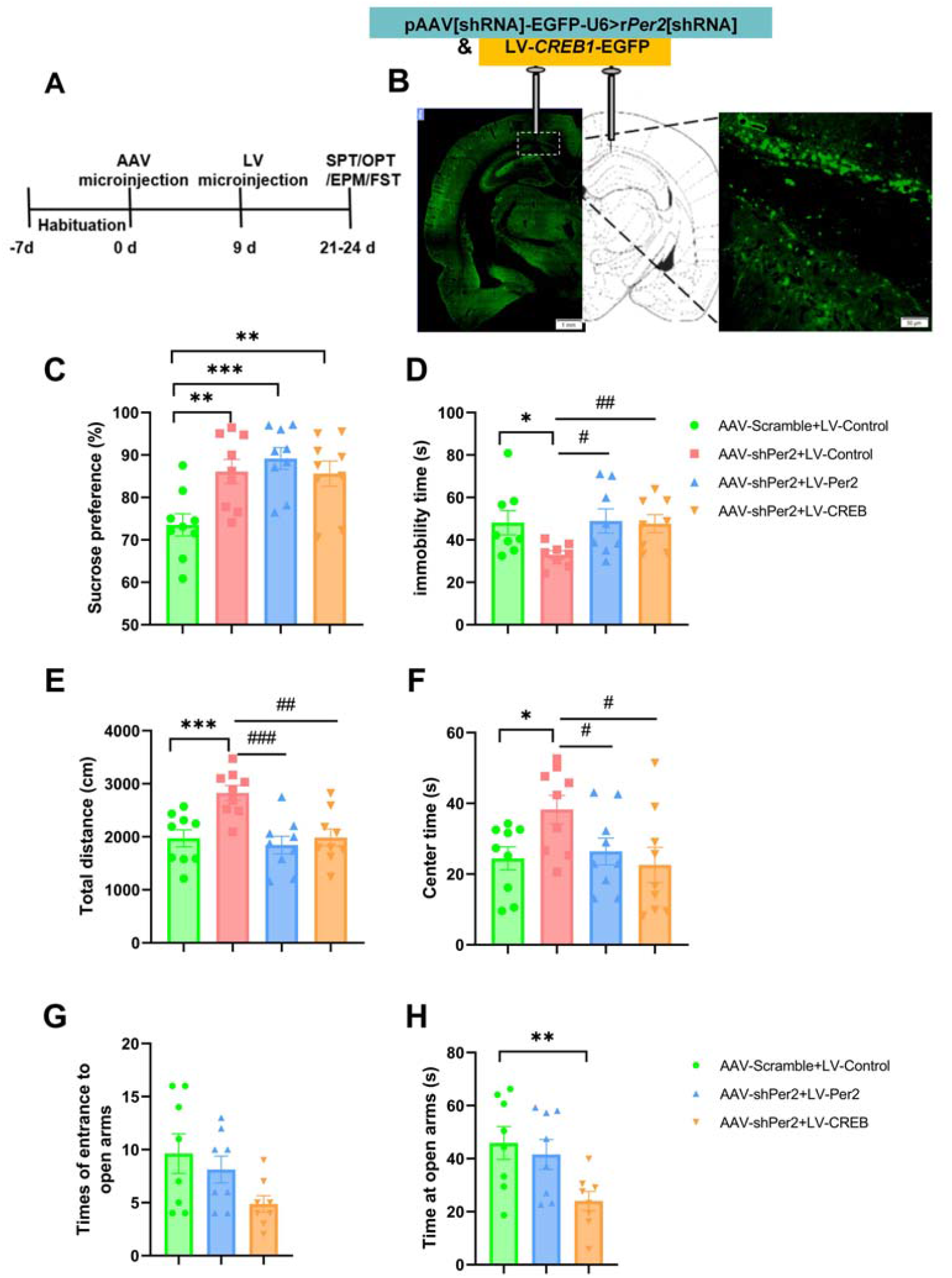
Knockdown of *Per2* in CA1 induced mania-like behaviors can be mitigated or converted by overexpression of either *Per2* or *CREB* in this region. (**A**) Timeline of this experiment. (**B**) Representative coronal section showing coinjection sites of AAV and LV (left panel, under 10 * object lens) and transfection cells (in fluorescence green, right panel, under 40 * object lens) in CA1 of the rat brain. Scale bars, 1 mm (left), 50 μm (right). (**C** and **D**) Data showing the effects of AAV-shPer2 and LV-Per2/CREB coinfused into CA1 on sucrose preference values in SPT (C) and immobility time in FST (D). Graphs are shown as means ± SEMs. *n* = 8-9. Values exceeded mean ± 2 * STD were excluded. \**P* < 0.05, \*\**P* < 0.01, \*\*\**P* < 0.001, vs. AAV-scramble + LV-control group; #*P* < 0.05, ##*P* < 0.01, vs. AAV-shPer2 + LV-control group; unpaired *t* test. \**P* < 0.05, #*P* < 0.05, Mann Whitney test (D). (**E** and **F**) Results of OPT: total distance (E) and time spent in central zones (F) in the four groups. Graphs are shown as means ± SEMs. *n* = 9. \**P* < 0.05, \*\*\**P* < 0.001, vs. AAV-scramble + LV-control group; #*P* < 0.05, ##*P* < 0.01, ###*P* < 0.001, vs. AAV-sh*Per2* + LV-control group; unpaired *t* test. (**G** and **H**) Results of EPM: times of entries to open arms (G) and time spent at open arms (H). Graphs are shown as means ± SEMs. *n* = 8. \*\**P* < 0.01, vs. AAV-scramble + LV-control group; unpaired *t* test.

We also harvested CA1 brain tissues on day 21 of the experiment to examine molecular changes (fig. S7A). Quantitative reverse transcriptase PCR showed that ΔCT values of *Per2* were significantly upregulated in the AAV-sh*Per2* group, meaning that AAV-sh*Per2* significantly down-regulated *Per2* mRNA levels, which could be reversed significantly by LV-*CREB* in CA1 (fig. S7C), demonstrating a positive regulated effect of CREB on *Per2* expression at transcriptional levels. Nevertheless, we didn’t find any significant difference in ΔCT values of *Creb1* among groups (fig. S7B).

## Discussion

We demonstrated that CREB positively regulates pCREB and PER2 (fig. S5, C and D; fig. S6, C and D; S7C), and that PER2 positively modulates CREB (fig. S2E, S3E and S5E) and pCREB (fig. S3D) in a feedback way, forming a loop in CA1. Furthermore, our findings suggest that this feedback loop mediates the transition between depressive and manic like behaviors, seen in the summary schematic (fig. S7D). Specifically, knockdown of *Per2* in CA1 produced mania-like behaviors (Fig. 2), which were ameliorated or converted by overexpression of *CREB1* or itself in this region (Fig. 6). Similar results were found in that of *CREB1*: knockdown of *CREB1* in CA1 produced mania-like behaviors, which could be improved by overexpression of *Per2* in this region (Fig. 4), and overexpression of *CREB1* in CA1 leads to the opposite effect, which could be reversed by knockdown of itself or *Per2* (Fig. 5). Importantly, our findings provide an insight into the molecular mechanisms underlying transition between manic and depressive episodes. Currently, there is no widely-accepted BD animal model exhibiting both depression and mania-like phenotypes (*17, 40, 41*). We present here a novel animal model exhibiting both characteristic phenotypes of BD, produced by regulating only one gene, avoiding drug-addicted effects of typical pharmacological models (*41*).

### Roles of CREB, pCREB and PER2 in mood regulation

Multiple studies have shown that CREB/pCREB/PER2 participate in the pathogenesis of BD (*21-23, 42, 43*). In this study, we found that lithium could rescue the ouabain-induced reductions in the levels of pCREB and PER2 in CA1, in line with previous reports (*22, 23*). We didn’t find any effect of lithium on CREB levels (Fig. 1F), probably due to that lithium robustly increases cAMP-induced CRE-mediated transcription, as well as CREB activity, the latter of which cannot be detected by western blot (*44*). It is reported that lithium may regulate *Per2* expression at the transcriptional level (*22, 38*), and that CREB is not essential for the effect of lithium on rhythm amplitude (*38*).

To the best of our knowledge, this may be the first report using knockdown and overexpression of *Per2* and *CREB1 in vivo* to reveal the pathogenesis of bipolar transformation. We propose that PER2 may be the core protein mediating mood transition, for the following reasons: i) CREB is a downstream effector of various intracellular signaling pathways, and therefore, can be affected by multiple factors (*27, 28, 45*); ii) during activation, CREB is phosphorylated into pCREB, which can translocate to the nucleus and regulate the expression of hundreds of genes (including c-*fos, Per1, BDNF* (*31*) and *Homer1*, thereby impacting a multitude of pathophysiological processes (*27, 28, 45*); iii) PER2 has not been reported to be involved in many pathways or physiological processes, indicating a relative specificity of its action in the pathophysiology of BD.

### CREB–pCREB–PER2 forms a novel feedback loop

As CREB is upstream to pCREB and *Per2*, the CREB–pCREB–PER2 pathway is already well-known. Intriguingly, we found that PER2 also positively regulates CREB, forming a positive feedback loop, which may be of significance to known molecular circadian feedback loops (*46*). As quantitative reverse transcriptase PCR results showed no significant effect of PER2 on *Creb1* mRNA levels (fig. S7), this feedback regulation cannot be deduced from transcriptional levels. A recent study by Brenna et al. showed that PER2 physically interacts with CREB and regulates its binding to the CRE element on *Per1* to induce expression (*47*). Therefore, it is intriguing whether PER2 also regulates its own expression via interaction with CREB and its co-regulator. Further study is needed to elucidate whether the PER proteins positively regulate CREB at the post-translational levels.

### Limitations of the study

This study was only from the perspective of cellular pathway and shows the non-circadian function of clock proteins. All brain tissue samples were collected consistently at one time point, 10:00 am, and therefore cannot reflect circadian fluctuations (*48*). Because reduced circadian amplitude may be a key feature of BD (*49, 50*), and lithium acts on *Per2* via enhancement of the baseline rhythm amplitude (*21, 38*), research into the expression levels of *CREB1* and *Per2* in this feedback pathway may be important in revealing the pathogenesis of BD. Thus, to investigate the rhythmicity of this loop, further research is needed to reveal the alterations in protein or mRNA levels in CA1 over a 24 h period. Moreover, we performed microinjections into the CA1 region and extracted proteins or mRNAs without cell-type specificity. Therefore, whether this loop has different functions in neurons and glial cells needs to be investigated in future studies.

In addition, we conducted this study in rats, and the findings therefore need to be confirmed by clinical studies, and proteins in the CREB–pCREB–PER2 feedback loop may have potential as state-dependent biomarkers in BD.

## Acknowledgements

I sincerely thank all of my supervisors, including Prof. Wei Chen during my master stage, Prof. George Fu Gao, Lin Lu and Su-Xia Li during my doctoral stage, for their supervision, guidance and concerns for me. As this research work is all done by myself alone, I thank those hardships I overcame in the process, giving me more courage and willpower. Specially, I appreciate greatly to my family for their support and those passed time I spent in Shandong University. I also thank Barry Patel, PhD, from Liwen Bianji (Edanz), for editing the English text of a draft of this manuscript.

## Funding

National Natural Science Foundation of China (NSFC82201682)

Natural Science Foundation of Shandong Province (ZR2021QH282)

Fundamental Research Funds of Shandong University (2020GN095)

## Author contributions

XLW conceived the project, designed and performed the experiments, analyzed the data, wrote the paper and acquired the funds all by herself.

## Competing interests

Authors declare that they have no competing interests.

## Data and materials availability

All data are available in the main text or the supplementary materials.

## Notes

### Competing Interest Statement

The authors have declared no competing interest.

### Summary of Updates

The pCREB-Per2 pathway has been identified as a feedback loop: CREB-pCREB-Per2, thus, all of the manuscript including figures and supplemental files were all revised.

