## Supplemental Figures for "CREB-pCREB-PER2 feedback loop mediates transition between mania and depression-like behaviors"

**The PDF file includes:**

Materials and Methods

Figs. S1 to S7

Table S1

References (*31*, *51*)

Materials and Methods

**Materials**

Rats

Male Sprague Dawley (SD) adult male rats (aged 8-10 weeks, 220-240 g) were bought from Ji’nan Pengyue Experimental Animal Breeding Company (No. SCXK-2019-0003, Ji’nan, China) and housed in groups of three under a 9 h/15 h light/dark cycle (8:00 light on and 17:00 light off) with free access to food and water. The room temperature was maintained 23 ± 1°C and humidity was 50 ± 5% controlled. All experiments were performed in accordance with the National Institutes of Health Guide for the Care and Use of Laboratory Animals (China) and were approved by the Shandong University School of Basic Medicine Ethics Committee (Ethical approval number: ECSBMSSDU2022-2-51).

**Methods**

Stereotaxic surgery and intracranial microinjections

After anaesthetizing rats by isoflurane gas, we eliminated the fur from each rat skull and placed them on the stereotaxic apparatus. First, we inserted a cannula guide (internal diameter, 0.45 mm; RWD, Cat# 62001) stereotaxically into CA1 region (anterior/posterior, -4.3 mm; medial/lateral, ± 2.0 mm; and dorsal/ventral, -2.0 mm). Then, we microinjected AAV-shPer2/shCREB (1 μL per side) or AAV-Scramble (1μL per side) bilaterally into CA1 with Hamilton syringes (Hamilton, USA, Cat# 80300) connected to injectors (Hamilton, USA, Cat# 62201) via the cannula guide to inject within 5 minutes at a rate of 1 μL/min and kept the injector in place for an additional 3 minutes to allow diffusion. The lentiviruses (LV-Per2, LV-CREB and LV-Control) were microinjected in the same way (1 μL per side). After surgery, rats were singly housed and received penicillin 20 million units intraperitoneally per day for 3 days to prevent infection.

After experiments, we performed immunofluorescence to validate viral injection site and expression.

Ouabain-induced mania model

Ouabain-induced mania model was conducted according to published protocols (*31*). Ouabain was dissolved in artificial cerebrospinal fluid (aCSF, 1 mmol/L, 5 μL) and injected into the intracerebroventricular (anterior/posterior, -0.9 mm; medial/lateral, + 1.5 mm; and dorsal/ventral, -3.3 mm) at 1 μL/min in rats. The stereotaxic surgery process was the same as stated above.

Drug infusion

Drug administration was performed as described before (*31*). After ouabain microinjection surgery, rats received 0.9% saline (vehicle) or lithium carbonate (20 mg/kg, 2 mL/kg) intraperitoneally once per day at 9:00 a.m. consecutively for 7 days. Lithium carbonate was dissolved in saline at a concentration of 2 mg/mL.

Behavioral test procedures below were based on previous studies in our lab (*51*).

Sucrose preference test (SPT)

Rats were singly caged and habituated to water or 1% sucrose solution for 48 hours before SPT. Bottles were interchanged with positions at the middle time of the habituation stage. SPT was performed after rats were deprived with food and water for 4 h. Rats were given two bottles, one is tap water (W) and the other is 1% sucrose solution (S) with the positions of two bottles interchanged half an hour later to avoid side bias, the consumption of sucrose solution and tap water in 1 h was quantified by measuring the weight of each bottle, the sucrose preference was calculated as [Weight S/(Wei S + Wei W)]*100.

Forced swimming test (FST)

Rats were put into a glass barrel (25 cm in diameter) filled with water (45 cm depth) at 23 ± 1 ℃ under bright light conditions and video recorded for 5 min. A black baffle was set between two cylinders to prevent mutual affection. Cumulative immobility time was measured with a stopwatch. Immobility was defined as no movement of body, tails and four limbs, compared with mobility, which was recognized as swimming or struggling behaviors. Prolonged immobility time of rats was seen as a behavior of despair, while a shortened immobility time was identified as a mania-like effect.

Open field test (OPT)

Rats were placed in a plexiglas open-field arena (42 cm * 42 cm * 42 cm) and locomotion was monitored and tracked using an automated system (SMART v2.5.21 System, Panlab, USA) for 5 min. The area was divided into 25 panes that were equally in size. The central 9 panes were defined as the central zones and other panes were defined as the corner. Cumulative time spent in central zones and total distance was recorded.

Elevated plus maze (EPM) test

The elevated plus maze consists of a central platform with two closed arms (50 cm in length, 10 cm in width, 40 cm in height with walls on both sides) and two open arms in the same sizes without walls on either side. The maze was placed 100 cm above the floor. A rat was placed in the crossed area of the platform facing an open arm and permitted to travel in the maze at liberty for 5 min. Time spent in the open arms and number of entrances into open arms were parameters analyzed by SMART v2.5.21 software (Panlab, USA). The platform was cleaned with 75% ethanol after each test.

Rat tissue collection

For quantitative analysis of molecules, experiments were designed in parallel to behavioral studies and rats were decapitated without experiencing behavioral tests to avoid their influence on molecular alterations in the rat brain. These rats were anesthetized and brains were collected and froze at -80 ℃ for preparation of western blotting or RNA analysis.

Western blotting

Tissues were homogenized in a RIPA lysis buffer (Macklin, cat#R917927-100ml) mixed with protease inhibitor cocktail (Sigma-Aldrich, cat#P8849-1ML) and protease phosphatase inhibitor mixture (Innochem, cat#B2248). Samples were then centrifuged for 30 min at 10,000 g at 4 ℃, and the supernatant was collected and quantified using the Microplate BCA Protein Assay Kit (Thermo Fisher Scientific). Equal amounts of protein sample were separated by 10% SDS-PAGE gels. Then proteins were transferred onto Hydrophobic PVDF Transfer Membrane (Millipore, cat# SJHVM4710). After that, membranes were washed by TBST (Tris-buffered saline with 0.1% tween 20), blocked by 5% BSA for 1 h and then incubated in primary antibody overnight at 4 ℃. Following primary antibodies were used in the study: mouse anti-GAPDH (1:5000, Bioss), mouse anti-Per2 (1:1000, Proteintech), rabbit anti-phospho-CREB (Ser133) (1:1,000, Cell Signaling Technology), rabbit anti-CREB1 (1:1000, Proteintech). The next day, after washed 5 min with TBST for 3 times, membranes were incubated with HRP conjugated secondary antibody (1:2,000, Bioss) for 1 h at room temperature, followed by 5 min washes for 3 times with TBST. Through drip with western chemilumi HRP substrate (Millipore, cat# WBKLS0100), the protein strips were imaged in the fluorescent imager (Tanon-5200). Protein levels were normalized to GAPDH on the same gel. All quantification was performed with Image J and Graphpad prism 8 software.

Quantitative reverse transcriptase PCR

The CA1 tissue was quickly harvested from the rat brains in a freezing microtome (Leica) and frozen in -80℃. The tissues were homogenized by a low-temperature beveller (Servicebio, KZ-Ⅲ-FP), and RNA was extracted by Trizol. cDNA was synthesized using RT First Strand cDNA Synthesis Kit (Servicebio, cat#G3337). Quantitative real-time PCR analysis was performed using 2×SYBR Green qPCR Master Mix (None ROX, Servicebio), following the standard protocol on the Real-Time PCR System platform (Bio-rad, cat#CFX). *GAPDH* was used as an internal control for normalization with the *Per2* and *Creb1*. The primers used were listed in Table S1.

Immunofluorescence

After behavioral experiments, to validate viral injection site and transfection results, we also anesthetized parts of those rats experienced behavioral tests and conducted intracardiac perfusion with phosphate-buffered saline (PBS) followed by 4% paraformaldehyde (PFA). Brains were postfixed in PFA for 24 h at 4 ℃, and then transferred to 30% sucrose in PBS to prepare for immunofluorescence. Coronal slices were cut on a freezing microtome (Leica, CM1950) at a 25-μm thickness. After repeated rinsing in PBS for 3 times with 5 min each, brain slices were hanged on the glass slide with antifade slide mounting medium (Solarbio). Images were photographed using a full field digital slice scan microscope (vs120, Olympus VS120, Japan) and analyzed using OlyVIA software (Olympus, Japan).

Statistical analysis

All data were analyzed using GraphPad Prism 8 and are shown as mean ± SEM. Animals with missed virus injection and values exceeded mean ± 2 * STD were excluded from analysis. The statistical tests used for the behavioral data and western blotting data were two-tailed Student’s *t* test or Mann Whitney test between different groups. Data of qPCR was analyzed by two-tailed Student’s *t* test. Data distribution was assumed to be normal when *t* tests were performed, while, when the normality test showed significant difference, Mann Whitney tests were used. *P* < 0.05 was defined as statistical significance.

Fig. S1.

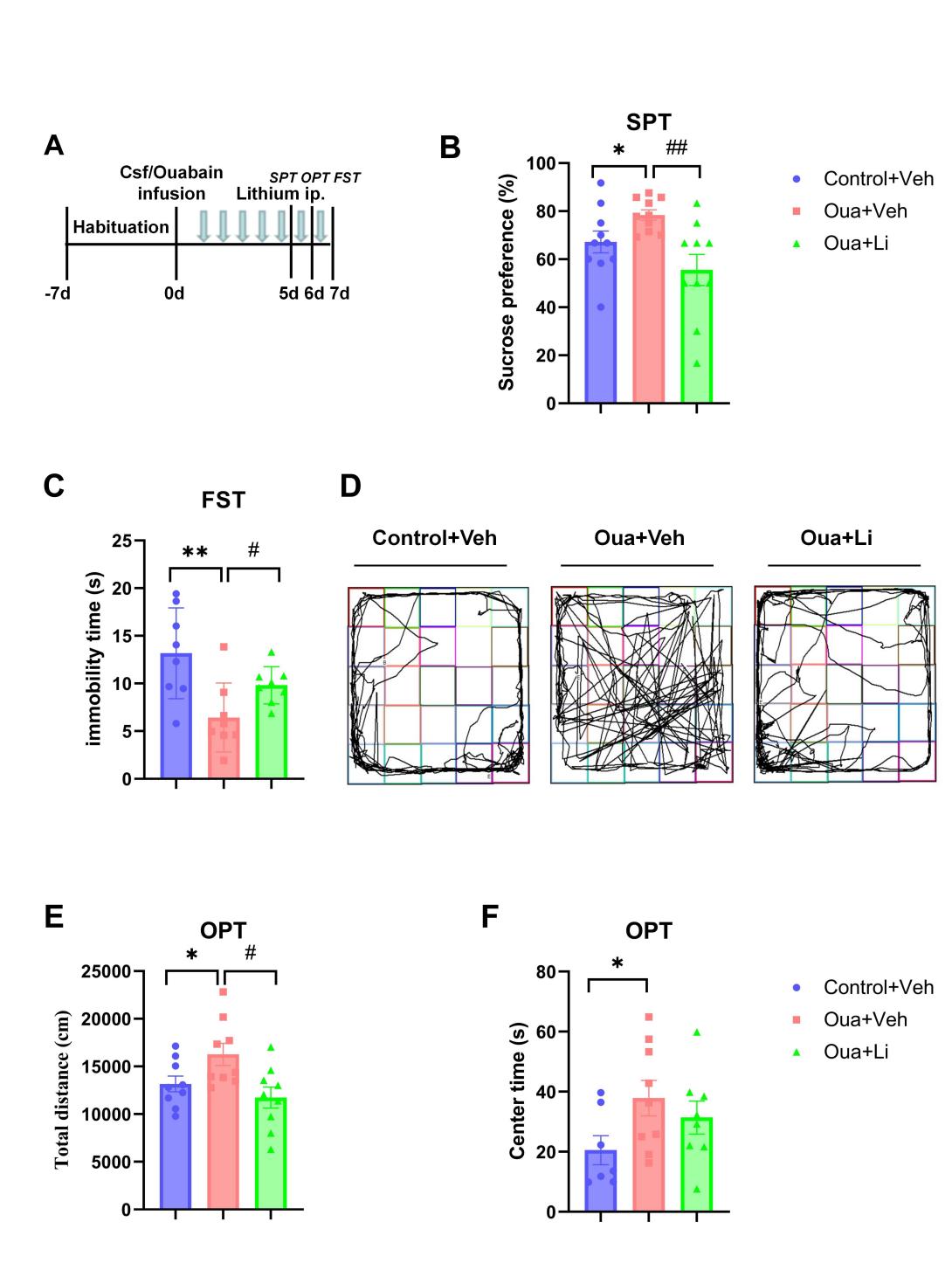

**Figure S1. Ouabain induces mania-like behaviors**

(A) Experimental timeline.

(B and C) Bat graphs show the effects of ouabain and lithium on sucrose preference values in SPT (B) and immobility time in FST(C). Data are shown as mean ± SEM. *n* = 8–10. **P* < 0.05, ***P* < 0.01, vs. Control+Vehicle group; #*P* < 0.05, vs. Ouabain + vehicle group. Unpaired *t* test. ##*P* < 0.01, vs. Ouabain + vehicle group, Mann Whitney test (B).

(D) Representative images of rat traces in the three groups in open field.

(E and F) Effects of ouabain and lithium on the total distance (E) and time in central zones (F) in OPT. Data are presented as mean ± SEM, *n* = 7–9. **P* < 0.05 vs.Control+Vehicle group; #*P* < 0.05 vs. Ouabain + vehicle group; two-tailed unpaired *t* test.

Fig. S2.

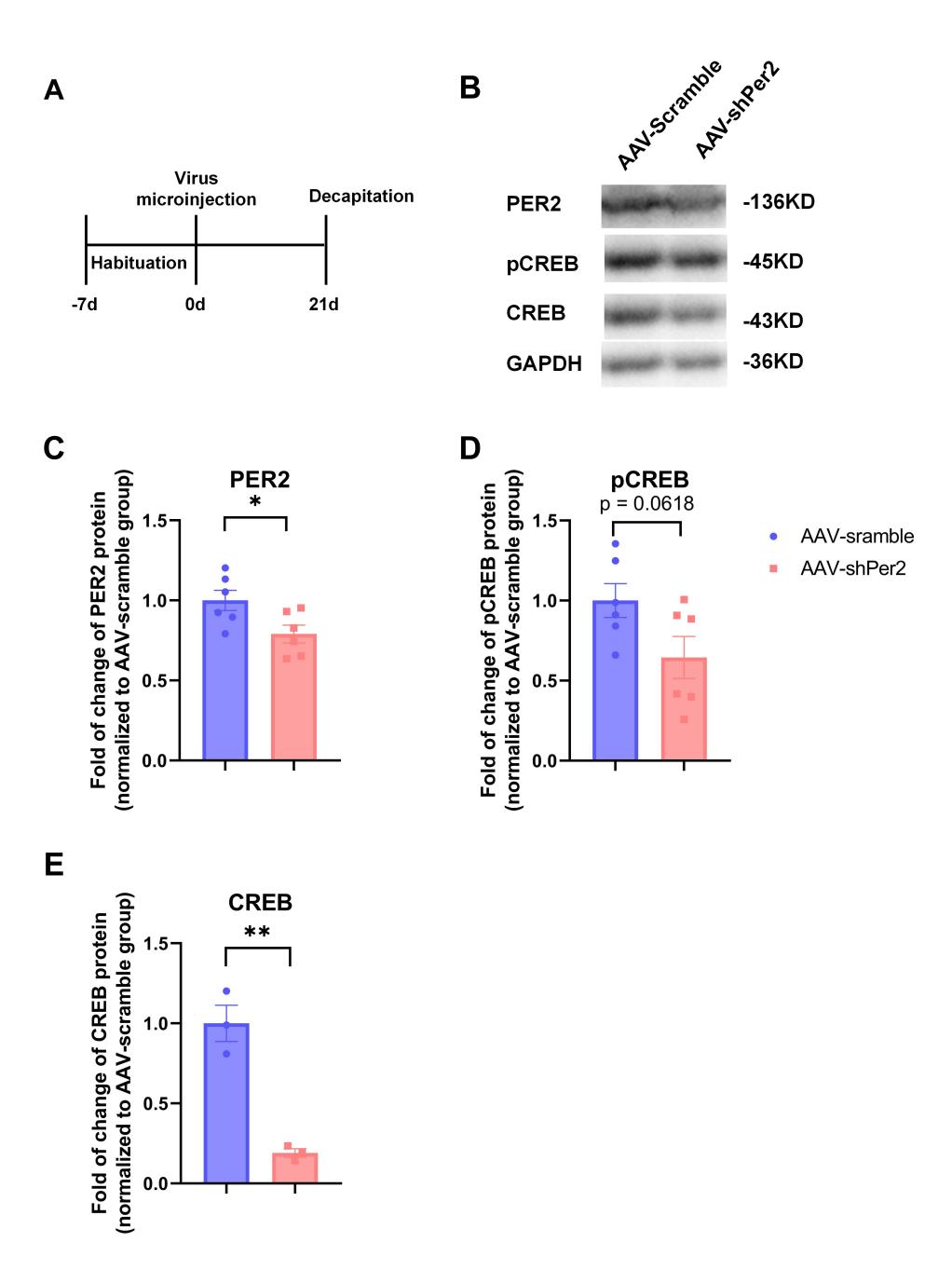

**Figure S2. Knockdown of *Per2* in CA1 downregulates CREB levels.**

(A) Timeline of the experiment.

(B) Representative bands of western blotting.

(C-E) Western blot analysis of PER2 (C), pCREB (D) and CREB (E). Graphs are shown as mean ± SEM. *n* = 3-6, values exceeded mean ± 2 * STD were excluded. **P* < 0.05; ***P* < 0.01; two-tailed unpaired *t* test.

Fig. S3.

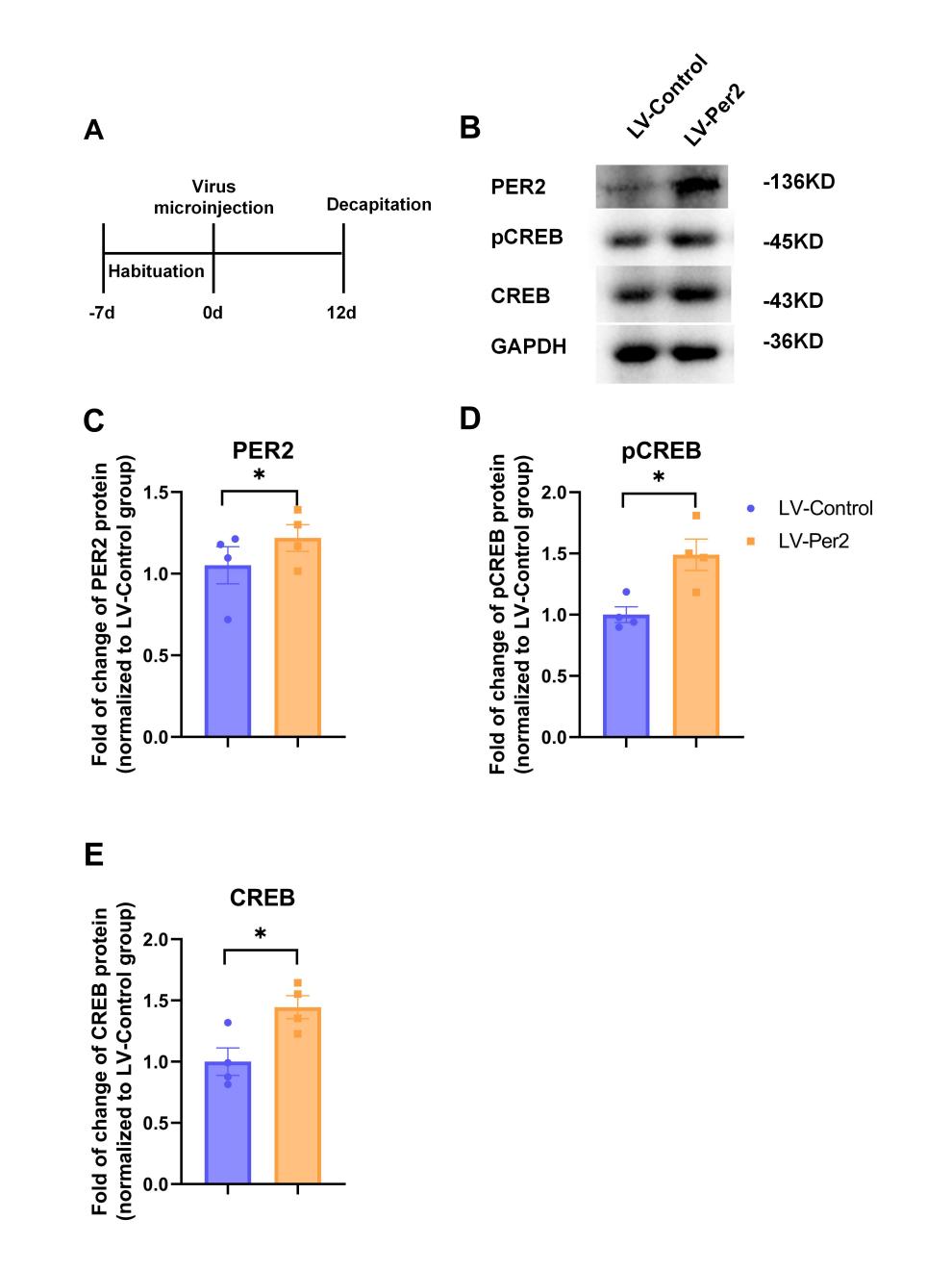

**Figure S3. Overexpression of *Per2* in CA1 upregulates CREB and pCREB levels**

(A) Timeline of the experiment.

(B) Representative bands of western blotting.

(C-E) Western blot analysis of PER2 (C), pCREB (D) and CREB (E). Graphs are shown as mean ± SEM. *n* = 4, values exceeded mean ± 2 * STD were excluded. **P* < 0.05, two-tailed unpaired *t* test.

Fig. S4.

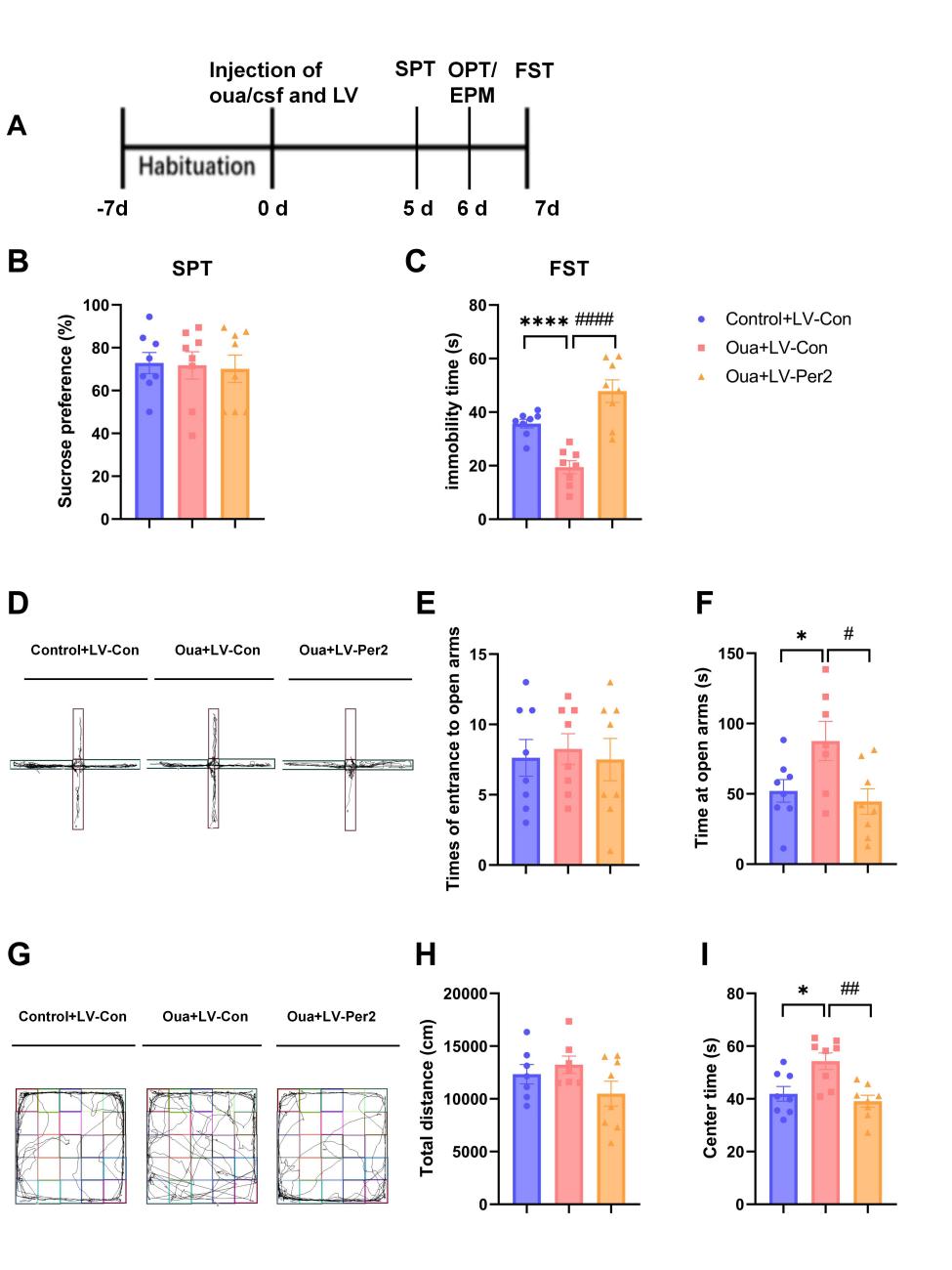

**Figure S4. Overexpression of *Per2* in CA1 region produces anti-mania-like effects**

(A) Timeline of the experiment.

(B and C) Bar graphs show the effects of LV-*Per2* and ouabain on sucrose preference values in SPT (B) and immobility time in FST (C).

(D) Representative images of rat traces of the three groups in elevated plus maze.

(E and F) Effects of LV-*Per2* and ouabain on times of entrance to open arms (E) and time at open arms (F).

(G) Representative images of rat traces of the three groups in open field test.

(H and I) Bar graphs show the effects of LV-*Per2* and ouabain on total distance (H) and time in central zones (I).

All data are shown as mean ± SEM. *n* = 7-8. Values exceeded mean ± 2 * STD were excluded. **P* < 0.05, *****P* < 0.0001, vs. Control + LV-control group; #*P* < 0.05, ##*P* < 0.01, ####*P* < 0.0001, vs. Ouabain + LV-control group; unpaired *t* test.

Fig. S5.

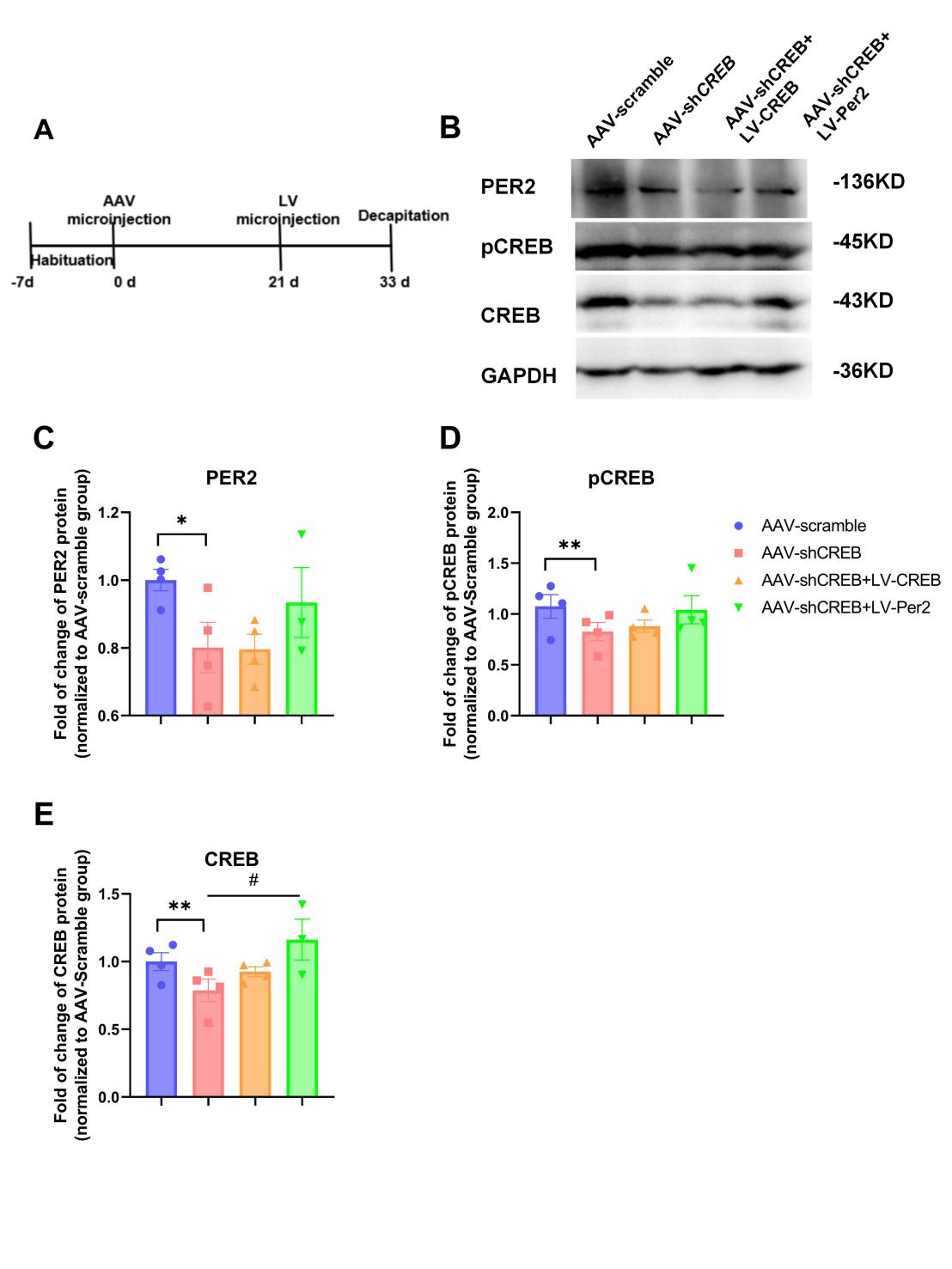

**Figure S5. CREB levels induced by knockdown of *CREB* can be rescued by overexpression of *Per2* in CA1**

(A) Timeline of the experiment.

(B) Representative bands of western blotting.

(C-E) Western blot analysis of PER2 (C), pCREB (D) and CREB (E) in CA1.

Graphs are shown as mean ± SEM. *n* = 3-4, values exceeded mean ± 2 * STD were excluded. **P* < 0.05, ***P* < 0.01 vs. AAV-scramble group; #*P* < 0.05, vs. AAV-shCREB group; two-tailed unpaired *t* test.

Fig. S6.

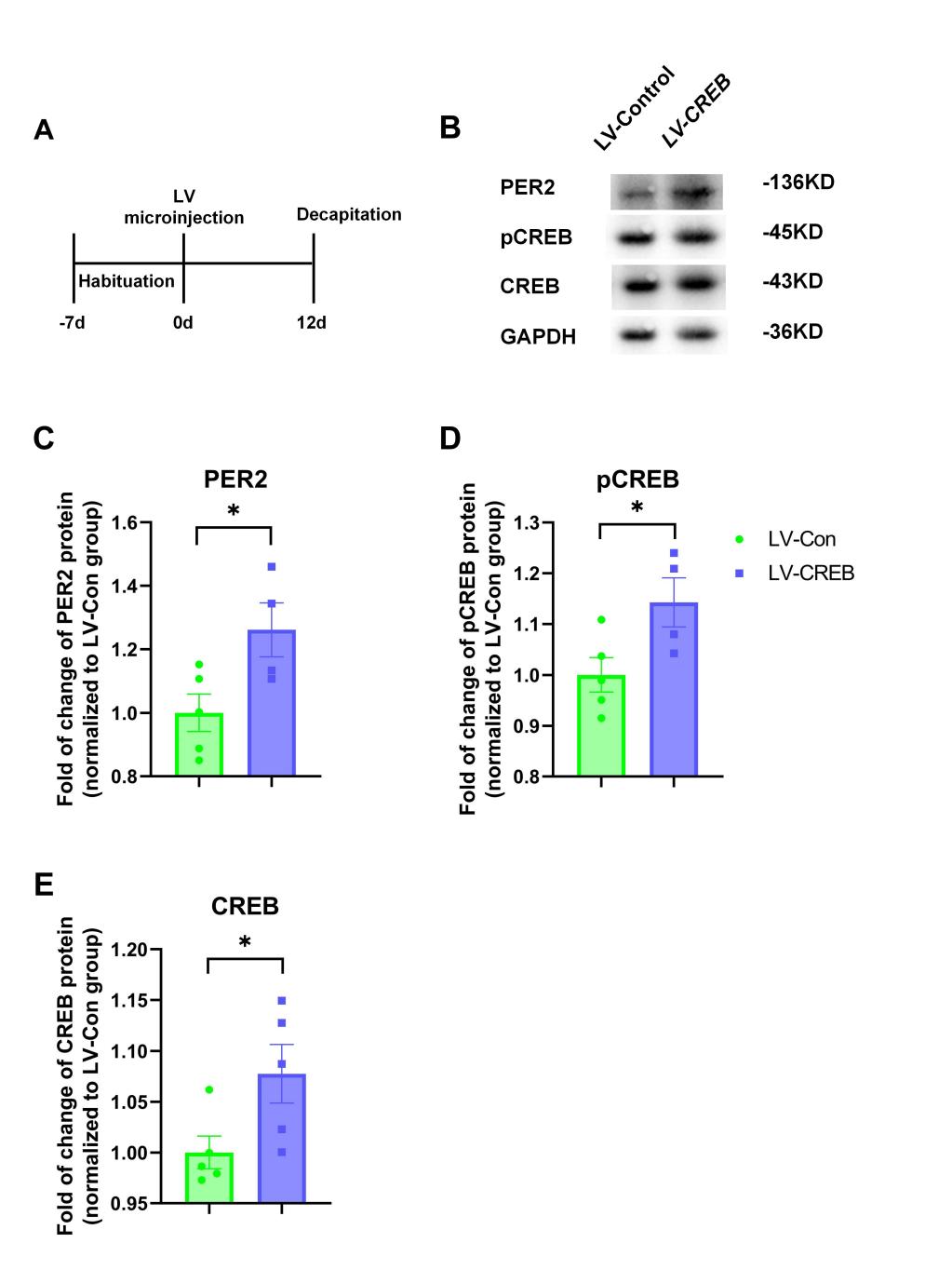

**Figure S6. Overexpression of *CREB* in CA1 upregulates pCREB and PER2 levels**

(A) Timeline of the experiment.

(B) Representative bands of western blotting.

(C-E) Quantification of the relative protein levels of PER2 (C), pCREB (D) and CREB (E).

All western blot data are shown as mean ± SEM. *n* = 4-5; Values exceeded mean ± 2 * STD were excluded. **P* < 0.05, Unpaired *t*- test.

Fig. S7.

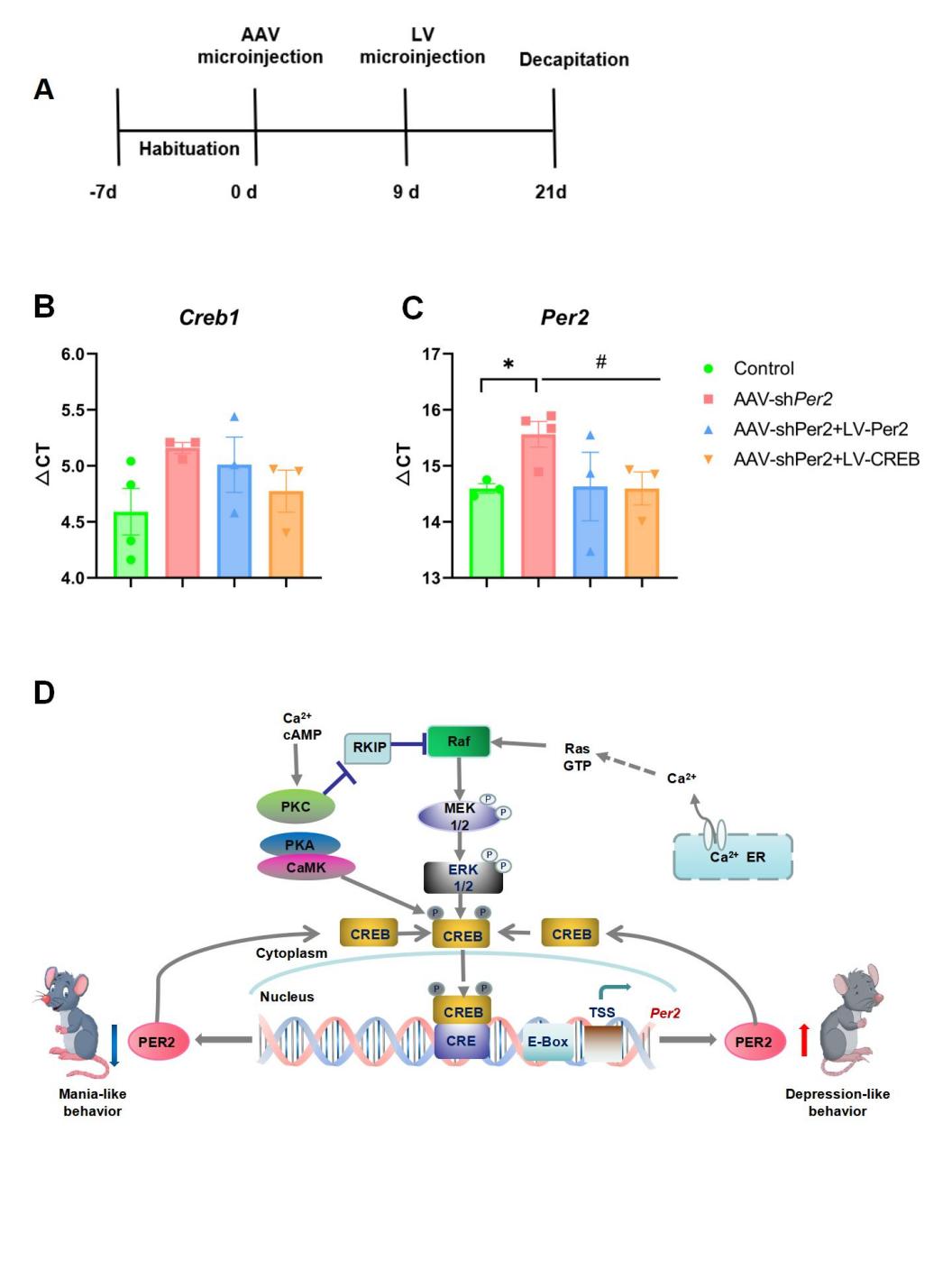

**Figure S7. Overexpression of *CREB* rescues *Per2* mRNA levels induced by knockdown of *Per2* in CA1 and schematic diagram**

(A) Timeline of the experiment.

(B-C) qRT-PCR shows the ∆CT values of *Creb1* and *Per2* genes in CA1. Data is shown as mean ± SEM. *n*=3-4; **P* < 0.05, vs. the control group;#*P* < 0.05, vs. the AAV-sh*Per2* group; unpaired *t* test.

(D) Schematic figure shows that the positive feedback loop of CREB-pCREB-PER2 in CA1 mediates transition between mania and depression-like behaviors. Upregulation of protein levels in this loop in CA1 produces depression-like behavior, while downregulation of those in this region leads to mania-like behavior in rats.

Table. S1. **Key resources table**

| REAGENT or RESOURCE | SOURCE | IDENTIFIER |
| --- | --- | --- |
| Antibodies | | |
| CREB1 Rabbit Polyclonal antibody | Proteintech | Cat#12208-1-AP; RRID:AB_2245417 |
| Phospho-CREB (Ser133)(87G3) Rabbit mAb | Cell Signaling Technology | Cat#9198; RRID:AB_2561044 |
| PER2 Mouse Monoclonal antibody | Proteintech | Cat#67513-1-Ig; RRID:AB_2882734 |
| Mouse Anti-GAPDH-Loading Control Monoclonal Antibody | Bioss | Cat#bsm-33033M |
| Bacterial and virus strains | | |
| AAV-shPer2 (pAAV[shRNA]-EGFP-U6>r*Per2*_shRNA) | Vector Builder | Cat#VB180626-1161npr |
| AAV-Scramble (pAAV[shRNA]-CMV-EGFP-U6>Scramble_shRNA) | Vector Builder | Cat#VB180117-1020znr |
| LV-Per2 (Ubi-*Per2*-EGFP-IRES-Puromycin) | Genechem | custom-made |
| LV-Control (Ubi-MCS-EGFP-IRES-Puromycin) | Genechem | Cat#LVCON136 |
| LV-CREB (Ubi-*Creb1*-EGFP-IRES-Puromycin) | Genechem | custom-made |
| AAV-shCREB (pAAV[shRNA]-EGFP-U6>r*Creb1*_shRNA) | Genechem | custom-made |
| Biological samples |  |  |
| Rat CA1 region brain tissue | This paper | N/A |
| Chemicals, peptides, and recombinant proteins | | |
| Ouabain | Sigma-Aldrich | Cat#4995-1GM |
| Lithium carbonate | Sigma-Aldrich | Cat#WXBD1461V |
| aCSF | Shanghai yuanye Bio-Technology | Cat#R22153-500ml |
| Experimental models: Organisms/strains | | |
| Rat: Crl:CD1(SD) | Ji’nan Pengyue Experimental Animal Breeding Company | 101 |
| Oligonucleotides | | |
| Primer: Creb1 Forward: TGGAGTTGTTATGGCGTCCTC | This paper | N/A |
| Primer: Creb1 Reverse:  AACCTCTCTCTTTCGTGCTGC | This paper | N/A |
| Primer: Per2 Forward:  TTACTGGTGGAAATAGCAAAGGCT | This paper | N/A |
| Primer: Per2 Reverse:  AGGGCCTTTTGTTTCCTCACTTT | This paper | N/A |
| Primer: GAPDH Forward: CTGGAGAAACCTGCCAAGTATG | This paper | N/A |
| Primer: GAPDH Reverse: GGTGGAAGAATGGGAGTTGCT | This paper | N/A |
| Recombinant DNA | | |
| shRNA targeting sequence: rPer2-RNAi:  CCACACTTGCCTCTGAAATAA | This paper | N/A |
| shRNA targeting sequence: rCreb1-RNAi: GCACTTAAGGACCTTTACTGC | This paper | N/A |
| Software and algorithms | | |
| Image J |  | N/A |
| GraphPad Prism | GraphPad Software | N/A |
| SMART v2.5.21 animal behavior analysis software | Panlab | N/A |

This table shows the detail information of key resources in this study.
